# Engineering a highly active thermophilic F_1_-ATPase by homolog-guided exploration and machine-learning-assisted prioritization

**DOI:** 10.64898/2026.08.27.747693

**Authors:** Ryohei Kobayashi, Kodai Miyake, Tatsuya Oya, Hiroshi Ueno, Yutaka Saito, Hiroyuki Noji

## Abstract

The rotary motor F_1_-ATPase has been extensively studied as a model molecular machine, yet rational engineering of its catalytic activity remains challenging because ATP hydrolysis is regulated by long-range intersubunit allostery and large conformational transitions. Here, we developed a homolog-guided engineering strategy to increase the maximum rotation rate of the thermophilic *Bacillus* PS3 F_1_-ATPase (TF_1_). Candidate mutation sites were first identified by comparing TF_1_ with the homologous enzymes bovine mitochondrial F_1_ (*b*MF_1_) and *Paracoccus* denitrificans F_1_ (PdF_1_), both of which exhibit higher maximum rotation rates than TF_1_. Systematic exploration of these sites identified four activity-enhancing hotspots, followed by focused hotspot exploration and machine-learning-assisted prioritization of combinatorial mutants. The best mutant, TF_1_(βY313L/βE332S), exhibited a 1.8-fold higher maximum rotation rate than TF_1_(WT) while retaining its functional thermostability. Interestingly, activity-enhancing substitutions were not limited to the residues conserved in both *b*MF_1_ and PdF_1_, indicating that the *b*MF_1_–PdF_1_ consensus substitutions effectively identify activity-enhancing hotspots rather than uniquely defining the optimal amino acid. Machine-learning-assisted exploration efficiently prioritized highly active mutants, although the predictive performance was limited by the relatively small training dataset and epistatic interactions among mutations. Kinetic and structural comparisons further provided mechanistic insights into the enhanced catalytic activity of the engineered mutant. Together, these results establish a practical strategy for engineering complex molecular motors by combining homolog-guided hotspot identification with focused hotspot exploration.

## Introduction

Biological molecular motors convert chemical energy into mechanical work and thereby drive diverse cellular processes, including intracellular transport, cell motility, and energy conversion (1, 2). Their high efficiency, nanoscale dimensions, and tunable mechanical outputs have made molecular motors attractive both as components of synthetic and biohybrid nanomachines and targets for protein engineering (3–5). Over the past two decades, molecular motors such as myosins, kinesins, and dyneins have been engineered to alter directionality, processivity, force production, and regulatory responses(6–9). By contrast, relatively few studies have attempted to increase the intrinsic speed or catalytic turnover of molecular motors (10, 11). Thus, whether highly evolved biological motors can be engineered for still higher performance remains largely underexplored.

F_1_-ATPase (F_1_), the soluble catalytic domain of F_o_F_1_-ATP synthase (F_o_F_1_), is one of the best-characterized molecular motors. In F_1_, ATP hydrolysis in the α_3_β_3_ stator ring is tightly coupled to unidirectional rotation of the central γ shaft (12–14). Decades of structural, biochemical, and single-molecule studies have established a detailed mechanochemical model of rotary catalysis, making F_1_ an ideal system for investigating the principles that govern molecular motor function. Despite this extensive mechanistic understanding, it remains unclear whether the rotational performance of F_1_ can be improved through protein engineering.

Among F_1_ homologs characterized to date, F_1_ from thermophilic *Bacillus* PS3 (TF_1_) has been a standard experimental model because of its high thermal stability, robust assembly, ease of handling, and suitability for single-molecule analysis. Single-molecule studies of TF_1_ established the canonical reaction scheme, in which each 120° rotation is coupled to the hydrolysis of one ATP molecule and is resolved into 80° and 40° substeps associated with ATP binding and ATP cleavage, respectively (15–18). The intervening waiting states are referred to as the binding dwell and catalytic dwell. Recently, cryo-EM studies have provided structural descriptions of these reaction intermediates, further supporting the mechanochemical model from single-molecule analyses (19, 20). These features have made TF_1_ a widely used platform for mechanistic studies and molecular engineering of rotary ATPases. However, TF_1_ rotates markedly more slowly than several mesophilic homologs, suggesting that robustness and rotation rates are not necessarily optimized simultaneously.

Comparative studies of F_1_ homologs have revealed substantial diversity in rotational rate. Under ATP-saturating conditions, TF_1_ has a maximum rate of approximately 180 rps, whereas *bovine* mitochondrial F_1_ (*b*MF_1_) rotates much faster, at approximately 707 rps (21). F_1_ from *Paracoccus denitrificans* (PdF_1_), which is proposed to share a common ancestor with mitochondrial F_1_, exhibits an intermediate maximum rotation rate of approximately 340 rps (22). Notably, the α, β, and γ subunits are highly conserved among these species, with particularly high sequence similarity in the β subunit. To identify the molecular determinants of these differences in rotational velocity, we previously analyzed hybrid F_1_s composed of subunits derived from TF_1_, *b*MF_1_, and PdF_1_ (23). This analysis showed that the maximum rotation rate is determined primarily by the β subunit, with a substantial contribution from α–β coupling. This conclusion is notable because the catalytic residues directly involved in ATP hydrolysis are completely conserved among these homologs and adopt nearly identical arrangements in available catalytic-dwell structures. Structural comparisons showed that active-site residues involved in ATP cleavage or phosphate release superpose with an RMSD of approximately 1 Å. These observations suggest that differences in rotation rates do not primarily arise from the canonical catalytic residues themselves, but from distal residues that influence conformational dynamics, stabilization of catalytic states, and inter-subunit communication. Such residues have remained comparatively underexplored as targets for improving the functions of molecular motor.

A major obstacle to engineering F_1_ has been the low throughput of conventional single-molecule analysis, which typically requires recombinant expression, purification, and characterization of individual mutants. We recently developed a high-throughput screening platform that integrates cell-free transcription–translation (TXTL) with multiplexed single-molecule rotation assays (24). This platform eliminates the need for cell culture and protein purification and enables rapid functional evaluation of large mutant libraries within a single experimental cycle. By combining a PURE-based cell-free expression system with single-molecule rotation measurements, the workflow from protein synthesis to functional characterization can be completed within approximately 10 hours. The platform therefore enables systematic exploration of sequence–function relationships in F_1_ at a scale that was previously impractical.

Here, we sought to improve the maximum rotation rate of TF_1_ by combining comparative sequence analysis, high-throughput cell-free single-molecule screening, and machine-learning (ML) assisted exploration (Fig. 1A). Guided by sequence comparisons with highly active mesophilic homologs, we systematically introduced substitutions into the α and β subunits and evaluated their effects using the TXTL-based screening platform (Fig. 1B). We further expanded the accessible sequence space using ML-guided prioritization of candidate mutations. Through this strategy, we identified TF_1_ mutants with substantially enhanced rotation rates while retaining the favorable characteristics of the thermophilic scaffold. Our findings demonstrate that TF_1_ is not a fully optimized endpoint but rather an engineerable molecular motor whose function can be further enhanced through protein engineering.

**Fig. 1.**
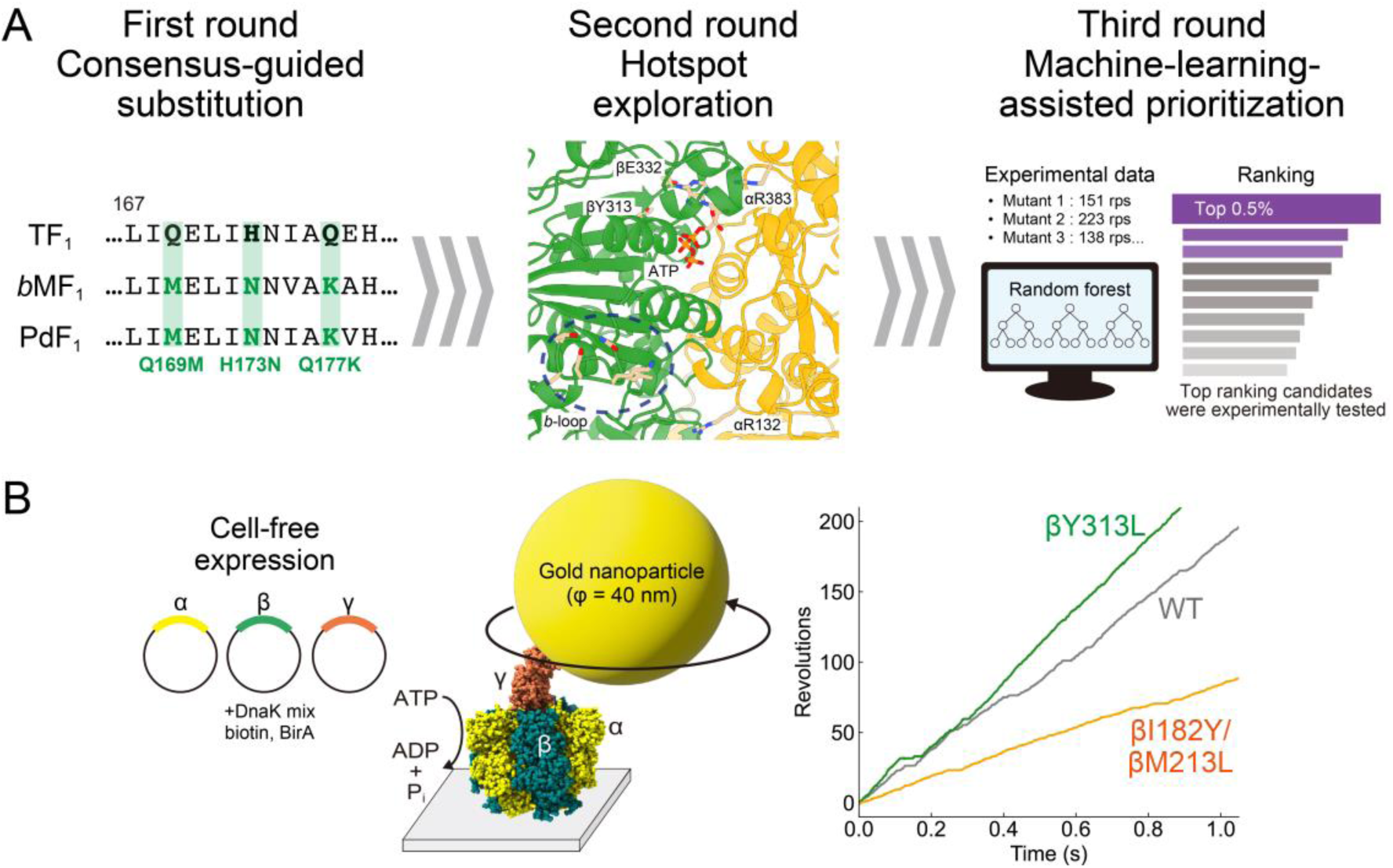
Overview of the engineering strategy. _(A)_ Experimental workflow for engineering TF_1_ with enhanced rotation rates through *b*MF_1_–PdF_1_ consensus substitutions, hotspot exploration, and machine-learning-assisted prioritization. (B) Schematic illustration of the cell-free single-molecule rotation assay. Representative rotation trajectories (right) were used to estimate the average rotation rates of individual mutants.

## Results

### Homolog-guided mutagenesis

Our previous work reported that the maximum rotation speed of F_1_ is primarily determined by the β subunit, although coupling between α and β also plays an important role (23). Based on this implication, we first compared the amino acid sequence of the β subunit from three representative F_1_ homologs with distinct maximum rotation rates: TF_1_, PdF_1_, and *b*MF_1_. We then identified candidate mutation sites where the TF_1_ residue differed from the residue conserved in both *b*MF_1_ and PdF_1_, hereafter referred to as the *b*MF_1_–PdF_1_ consensus residue. We excluded the N-terminal domain (1– 149 in TF_1_ numbering), which is essential for formation of the α_3_β_3_ hexameric ring (25), and the C-terminal domain (414–473), which is not directly involved in ATP hydrolysis, although it undergoes large conformational changes during rotation. Consequently, we focused on the central catalytic domain of the β subunit (150–413), which contains the P-loop (G159–T165), the catalytic glutamate (E190), and the DELSEED region (D390-D396) (Fig. 2A and Fig. S1). Within this region, 30 candidate sites were identified where the TF_1_ residue differed from the corresponding *b*MF_1_–PdF_1_ consensus residue (Fig. 2B and Fig. S2A), and these consensus residues were selected for substitution in the first-round mutagenesis (Fig. 2B). In addition to these simple substitutions, we also found the backbone difference between TF_1_ and *b*MF_1_-PdF_1_. A few residues are inserted in *b*MF_1_ and PdF_1_ to form a flexible loop (Fig. S1 and Fig. S2B).

**Fig. 2.**
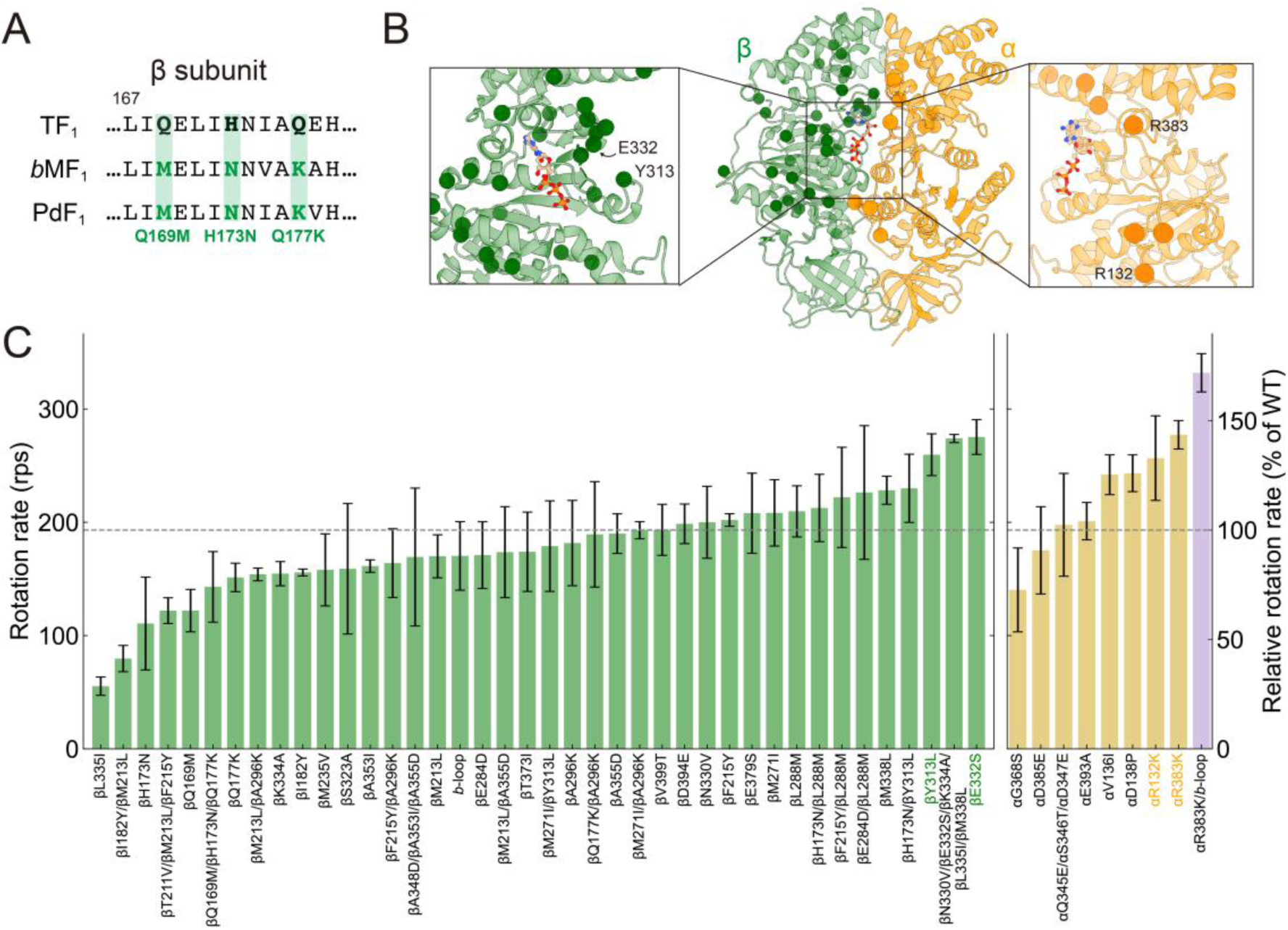
Homolog-guided mutagenesis by the first-round screen. (A) Partial sequence alignment of the β-subunits from TF_1_, *b*MF_1_, and PdF_1_. Residues conserved in *b*MF_1_ and PdF_1_ but differing in TF_1_ were selected as candidate substitution sites in TF_1_. Complete sequence alignments are shown in Figs. S1 and S3. (B) Candidate mutation sites mapped onto the αβ_DP_ interface of the TF_1_ structure (PDB: 7L1R). Candidate sites in the β- and α-subunits are shown as green and orange spheres, respectively. Enlarged views highlight Y313 and E332 in the β-subunit (left), and R132 and R383 in the α-subunit (right). (C) Average rotation rates of TF_1_ mutants carrying *b*MF_1_–PdF_1_ substitutions. Green and yellow bars indicate substitutions in the β- and α-subunits, respectively, and the purple bar indicates the αR383K/*b*-loop combination mutant. The right y-axis shows rotation rates relative to TF_1_(WT) (193 rps). Bars and error bars represent the mean ± SD for at least three independent molecules.

We first purified the two TF_1_ mutants: one containing all 30 *b*MF_1_–PdF_1_ consensus substitutions in the selected β-subunit region (referred to as TF_1_(β^30^)), and the other containing the loop insertion (Fig. S1 and Fig. S2B) corresponding to that found in *b*MF_1_ (referred to as TF_1_(*b*-loop)), respectively. Single-molecule rotation assays showed that both mutants rotated continuously in the counterclockwise direction with three distinct pauses at 3 mM ATP, similar to TF_1_(WT) (Fig. S2C). However, TF_1_(*b*-loop) did not improve the maximum rotation rate, whereas TF_1_(β^30^) exhibited a substantially lower rotation rate than TF_1_(WT) (Fig. S2D). These results suggested that simultaneous introduction of a large number of substitutions was not an effective strategy, probably because of unexpected allosteric effects among the substituted residues (26).

We therefore shifted our strategy to systematic evaluation of individual substitutions and small combinations of mutations. Because the candidate sites were distributed over a broad region of the β subunit, substitutions and their combinations were prioritized based on the feasibility of PCR-based plasmid construction. To evaluate a large number of mutants efficiently, we employed our previously developed high-throughput rotation assay based on a cell-free protein synthesis system (22) (Fig. 1B). Instead of the γ subunit containing two cysteine residues used in the conventional assay, we employed a γ subunit carrying two Avi-tags (referred to as TF_1_(WT) hereafter). Using this system, single-molecule rotation assays were performed at 3 mM ATP, and average rotation rates were estimated from at least three molecules (see *Methods* for details). Transient pauses arising from the ADP-inhibited state were excluded from the analysis because they reduce the time-averaged rotation rate and interfere with accurate estimation of the maximum rotation rate.

Fig. 2C summarizes the results of this first-round screening. Several substitutions increased the rotation rate relative to TF_1_(WT) (193 rps). Among the β-subunit mutants, βY313L (260 rps) and βE332S (275 rps) showed the largest improvements. Interestingly, a quintuple mutant carrying βN330V, βE332S, βK334A, βL335I, and βM338L rotated at a rate (274 rps) comparable to that of βE332S alone, although the single substitution βL335I markedly reduced the rotation rate (55 rps). These results demonstrate that homolog-guided substitutions can successfully identify activity-enhancing mutations, even in a complex molecular motor.

Previous experiments using hybrid F_1_s also suggested that interactions between the α- and the β-subunit contribute to the maximum rotation rate. We therefore expanded our analysis to the α-subunit by examining residues located at the α–β interface. Based on structural analysis of the α_DP_β_DP_ interface in the cryo-EM structure of TF_1_ (PDB: 7L1R), we selected ten additional candidate sites where TF_1_ residues differed from the corresponding *b*MF_1_–PdF_1_ consensus residues (Fig. 2B and Fig. S3). Single-molecule rotation assays identified four substitutions (αR383K, αR132K, αD138P, αV136I) that increased the rotation rate by more than 20% relative to TF_1_(WT), confirming the importance of α–β interactions in determining the rotation rate. Among them, αR383K exhibited the highest rotation rate (277 rps), followed by αR132K (257 rps). We also examined a preliminary combination of αR383K with the *b*-loop insertion because both are located at the α–β interface. Interestingly, this combination further increased the rotation rate to 332 rps, approximately 20% higher than αR383K alone, suggesting that the *b*-loop itself also contributes to activity enhancement. These hotspot positions were subsequently selected for comprehensive exploration in the second-round screen.

### Hotspot exploration

For further exploration, we selected hotspot sites where substitutions to the *b*MF_1_-PdF_1_ consensus residues enhanced activity in the first-round screen: βY313, βE332, αR132, and αR383 (Fig. 3A). We also included the *b*-loop because it further increased the rotation rate when combined with αR383K. To explore these hotspots more comprehensively, we examined alternative substitutions at each hotspot together with combinations of these hotspot mutations. The results are summarized in Fig. 3B. Notably, activity-enhancing substitutions were not limited to the *b*MF_1_-PdF_1_ consensus residues. A representative example is βY313: βY313W (275 rps) and βY313V (247 rps) increased the rotation rate to a similar extent as the consensus substitution βY313L, while little improvement was observed for βY313H (198 rps). These results suggest that some of the selected positions are potential catalysis-enhancing hotspots. Among the mutants examined in this second-round screen, the βY313L/βE332S double mutant showed the highest rotation rate (347 rps). The molecular basis of this enhancement is discussed later.

**Fig. 3.**
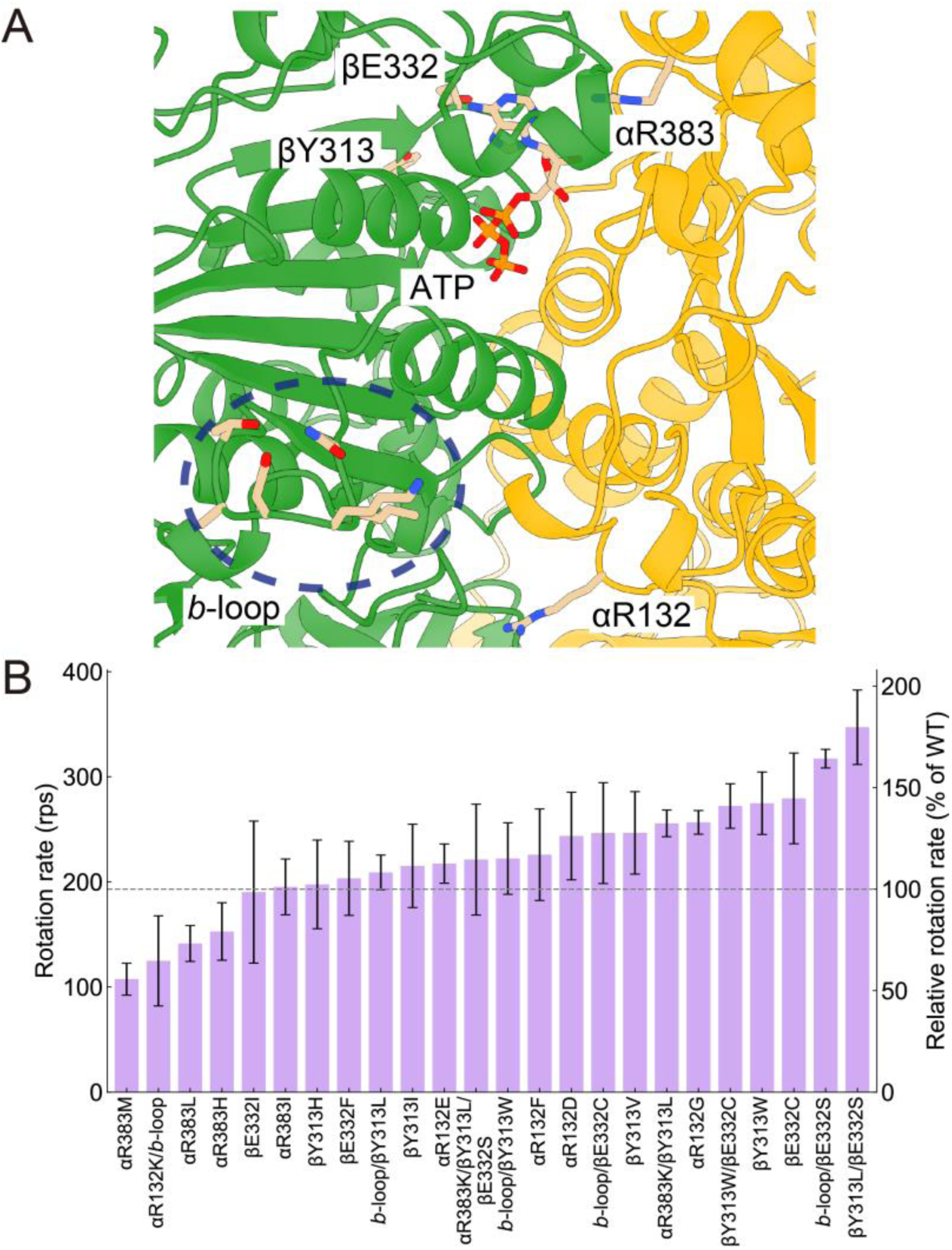
Hotspot exploration by the second-round screen. (A) Positions of βY313, βE332, αR132, αR383, and the *b*-loop mapped onto the αβ interface of an AlphaFold3-predicted structure. The β- and α-subunits are shown in green and yellow, respectively. ATP is shown in stick, and the *b*-loop is indicated by a dashed line. (B) Average rotation rates of TF₁ mutants tested in the second-round screening. Mutants include alternative substitutions at the identified hotspot positions and their combinations. The right y-axis shows rotation rates relative to TF_1_(WT) (193 rps). Bars and error bars represent the mean ± SD for at least three independent molecules.

Based on these experimental results, we next sought to identify mutants with even higher rotation rates by exploring combinatorial mutations across these hotspots. However, considering 20 possible amino acid substitutions at four distinct residue sites, together with the option of including the *b*-loop insertion, the total number of possible combinations was 320,000 (20⁴ × 2), which was impractical to test exhaustively with our current experimental setup. We therefore built a Random Forest regression model (see *Methods*) to prioritize candidates for experimental validation (Fig. 4A and Figs. S4-S6) (27, 28). The model was trained using the experimentally measured rotation rates of 30 mutants containing substitutions at βY313, βE332, αR132, αR383, and/or the *b*-loop insertion. The model showed moderate rank-ordering performance in five-fold cross-validation (Spearman’s ρ = 0.500, Fig. 4B), suggesting that the predicted rotation rates would help prioritize highly active candidates.

**Fig. 4.**
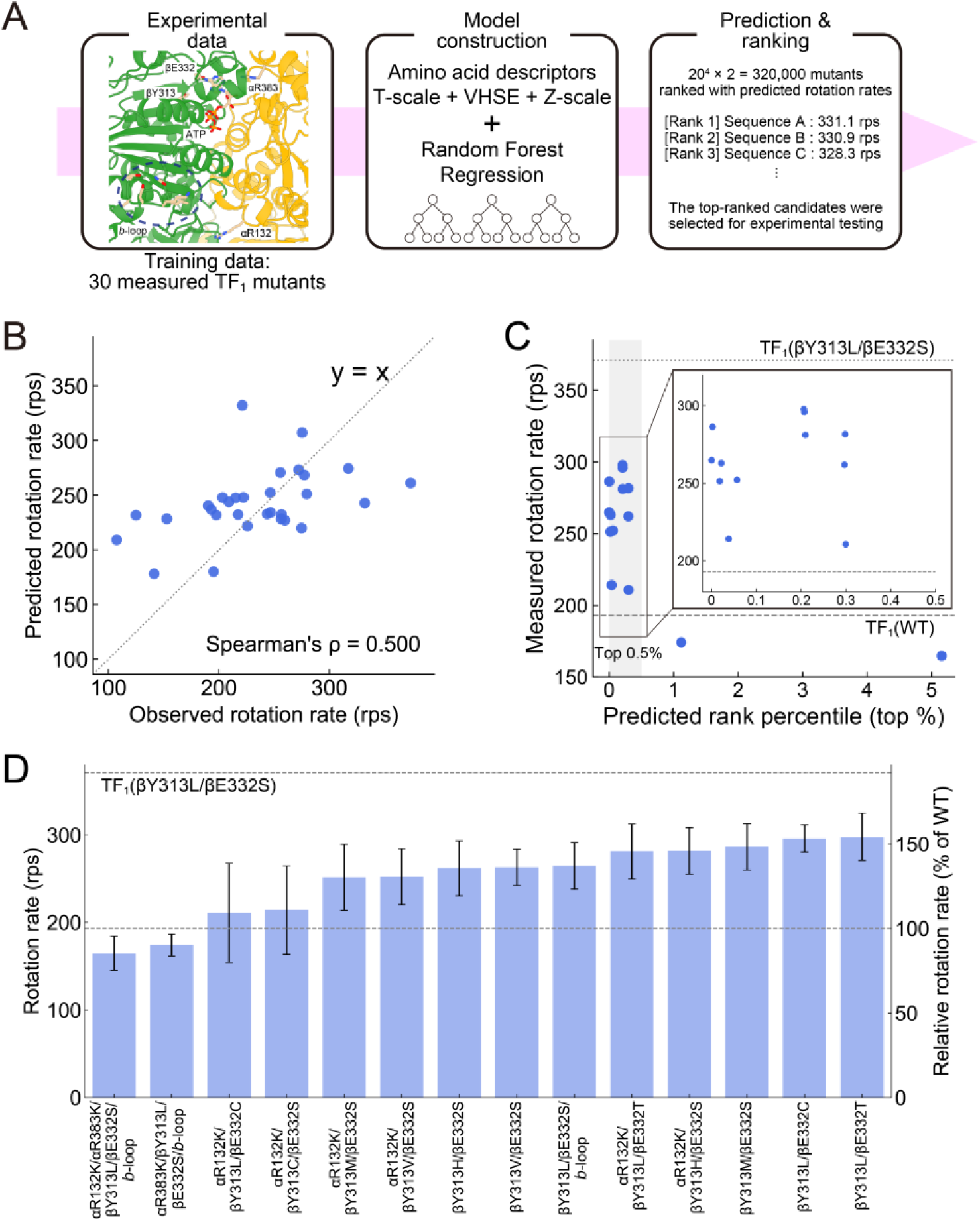
Machine-learning-assisted exploration by the third-round screen. (A) Workflow of machine-learning-assisted exploration. Rotation rates of 30 mutants carrying substitutions at βY313, βE332, αR132, αR383, with or without the *b*-loop insertion, were used to train a Random Forest model. The trained model was then used to predict and rank all 320,000 (20^4^ × 2) possible mutants, from which top-ranked candidates were tested experimentally. (B) Performance of the Random Forest model evaluated by five-fold cross-validation. Each point represents one experimentally characterized mutant. The x- and y-axes show the observed and out-of-fold predicted rotation rates, respectively. The dotted line indicates y = x. Spearman’s rank correlation coefficient was *ρ*=0.500. (C) Ranking of the experimentally characterized mutants among all 320,000 predicted mutants. Most mutants were ranked within the top 0.5% of the predicted mutants, whereas two mutants were ranked within the top 1% and top 5%, respectively. (D) Experimental validation of newly selected top-ranked candidates. The right y-axis shows rotation rates relative to TF_1_(WT) (193 rps). Bars and error bars represent the mean ± SD for at least three independent molecules.

We next selected previously untested, top-ranked candidates for experimental validation across the predicted landscape of 320,000 possible mutants. Most of the tested candidates ranked within the top 0.5%, although two lower-ranked candidates from the top 1% and top 5% were also included (Fig. 4C). The tested candidates within top 0.5% exhibited higher rotation rates than TF_1_(WT), indicating that the model effectively enriched highly active mutants, although it did not accurately predict absolute rotation rates (Fig. S6). Consistent with the moderate rank-ordering performance observed in cross-validation, the two lower-ranked candidates showed lower rotation rates than those within the top 0.5% candidates. However, contrary to our expectations, none of the newly tested candidates exceeded TF_1_(βY313L/βE332S), the best mutant identified in the second-round screen (Fig. 4D). Notably, the quintuple mutant containing αR132K, αR383K, βY313L, βE332S, and the *b*-loop insertion showed a lower rotation rate than TF_1_(WT), reaching only 165 rps. These results indicate that individually beneficial mutations are not necessarily additive when combined, suggesting that epistatic interactions or long-range allosteric constraints restrict the sequence space for achieving faster rotary catalysis.

### Thermostability analysis of the best mutant

Finally, we examined the thermal stability of TF_1_(βY313L/βE332S), hereafter referred to as TF_1_(LS), the fastest TF_1_ mutant identified in this study, together with TF_1_(WT) and *b*MF_1_(WT) for comparison (Fig. 5). Thermal stability was evaluated by comparing residual ATPase activity after heat treatment.

**Fig. 5.**
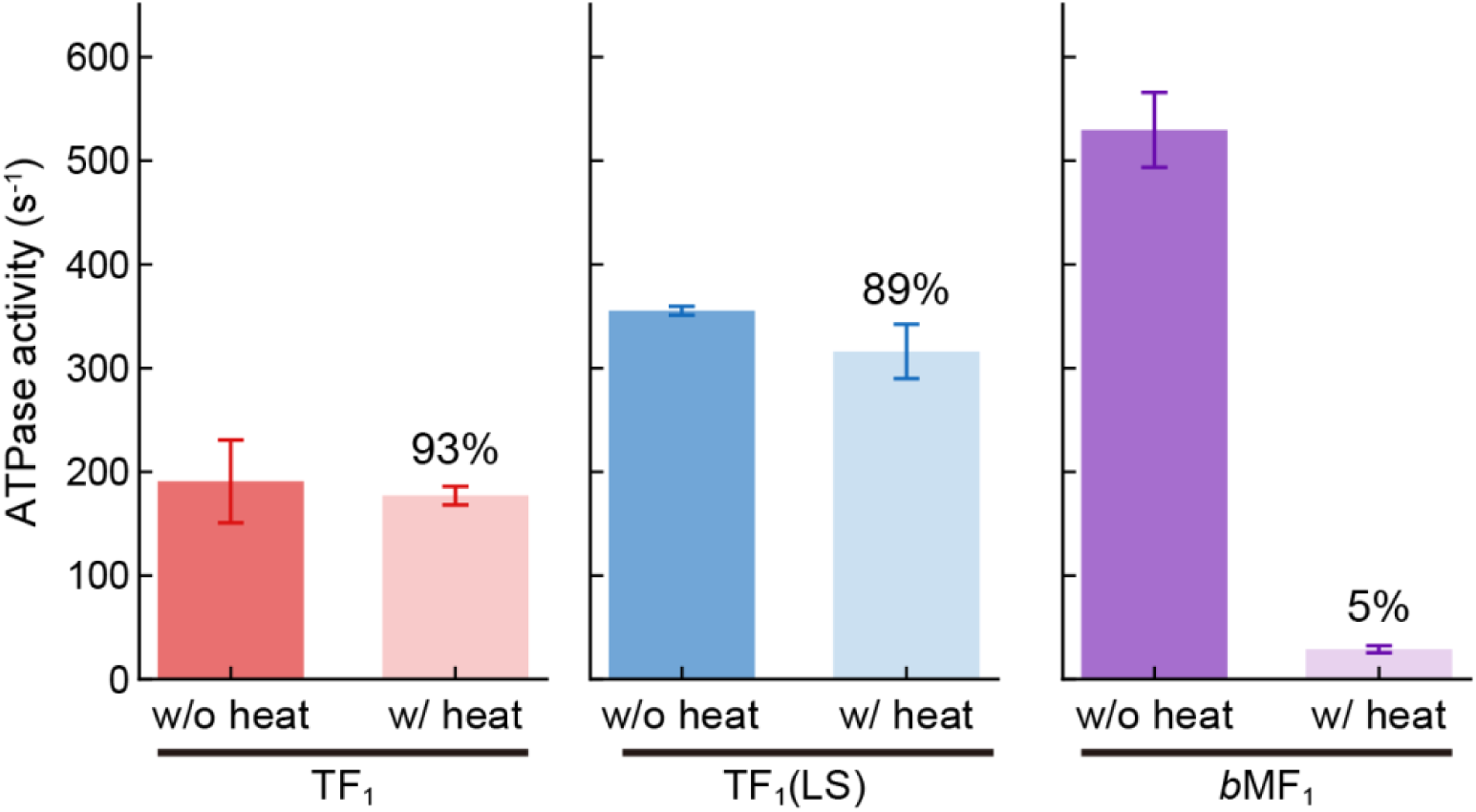
Functional thermal stability of the best mutant. ATPase activity before (left) and after (right) heat treatment is shown in each panel. Bars and error bars represent the mean and SD from three independent experiments. Percentages shown in each panel indicate the relative activity after heat treatment, with the untreated sample set to 100%.

Before heat treatment, ATPase activities followed the same trend as the single-molecule rotation assay: *b*MF_1_(WT) showed the highest activity, followed by TF_1_(LS), whereas TF_1_(WT) exhibited the lowest activity. For all three enzymes, ATPase activities measured in solution were lower than the ATP hydrolysis rates estimated from the maximum rotation rates in the single-molecule assay. This difference is commonly observed in F_1_ and is mainly attributed to ADP inhibition in the biochemical assay (29). Following heat treatment at 65°C for 10 min, *b*MF_1_(WT) retained only approximately 5% of its initial ATPase activity. In contrast, both TF_1_(WT) and TF_1_(LS) largely maintained their ATPase activities. These results demonstrate that TF_1_(LS) retained the functional thermal stability of TF_1_(WT) while exhibiting enhanced catalytic activity.

### Kinetic characterization of the best mutant

To investigate the kinetic basis of the enhanced rotation rate of TF_1_(LS), we performed single-molecule rotation assays over a wide range of ATP concentrations using purified F_1_ molecules. TF_1_(WT) was analyzed in parallel for comparison (Fig. 6A). Michaelis–Menten analysis showed that TF_1_(LS) exhibited an approximately two-fold higher maximum rotation rate 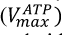 than TF_1_(WT), whereas the ATP-binding rate constant estimated from 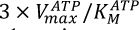 was nearly identical between the two enzymes. These results indicate that the enhanced rotation rate of TF_1_(LS) does not arise from accelerated ATP binding but from an increase in the turnover rate after ATP binding.

**Fig. 6.**
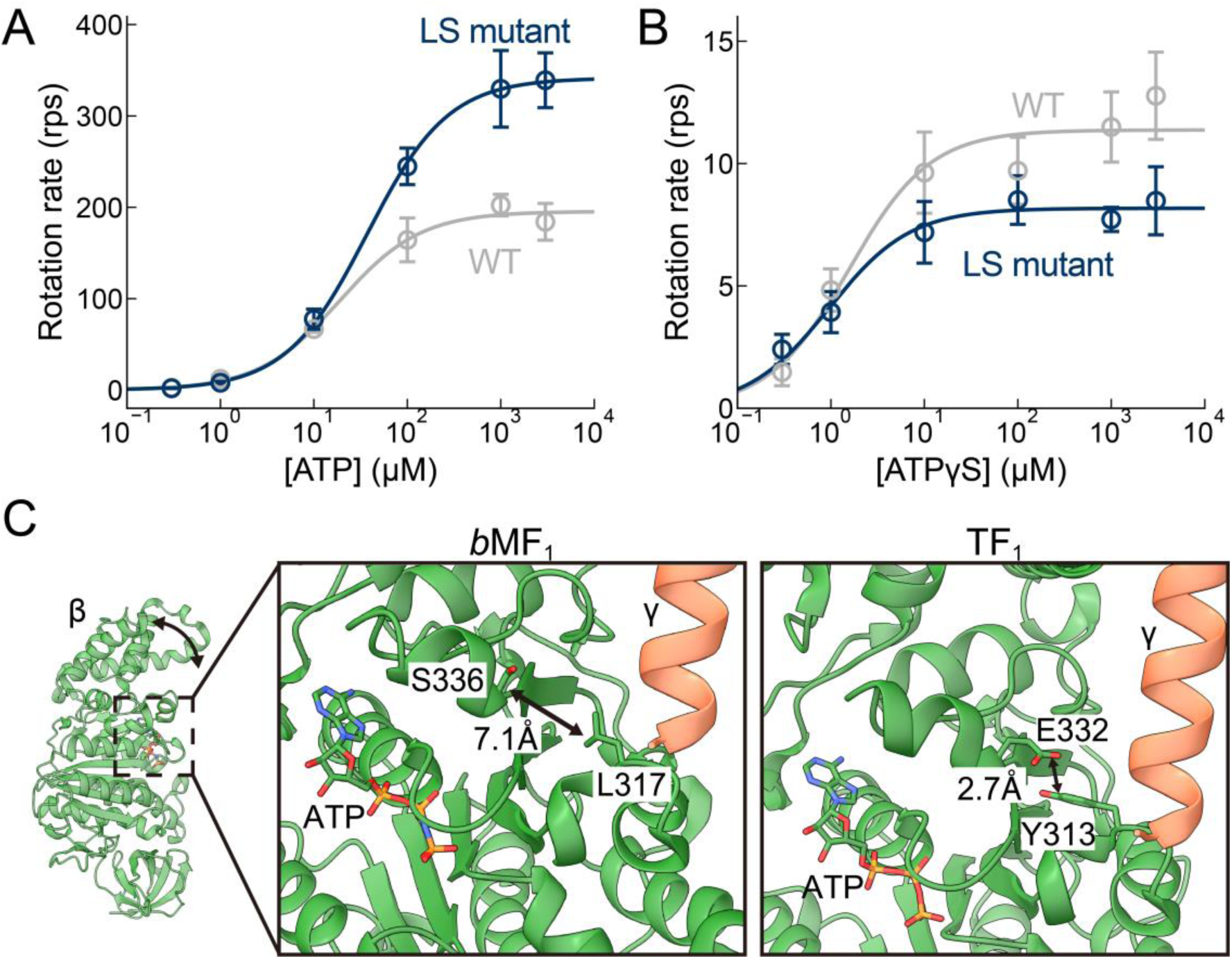
Kinetic and structural comparison of the LS mutant. (A) Michaelis–Menten analysis of ATP-driven rotation in TF_1_(WT) (gray) and TF_1_(βY313L/βE332S), termed the LS mutant (blue). Circles and error bars represent the mean value and SD, respectively (n = 5-7 molecules for each datapoint). Curves represent fits to the Michaelis–Menten equation, 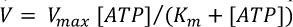 giving the following fitted parameters with errors estimated from the fit. 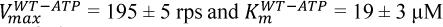 for TF_1_(WT), and 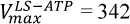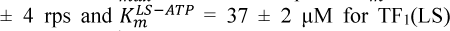. The binding rate constant estimated from 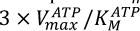 were 3.1 × 10^7^ M^-1^ s^-1^ and 2.8 × 10^7^ M^-1^ s^-1^ for TF_1_(WT) and TF_1_(LS), respectively. (B) Michaelis–Menten analysis of ATPγS-driven rotation in TF_1_(WT) (gray) and TF_1_(LS). Circles and error bars represent the mean value and SD, respectively (n = 5-10 molecules for each datapoint). Curves represent fits to the Michaelis–Menten equation, 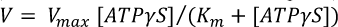, giving the following fitted parameters with errors estimated from the fit. 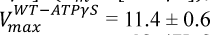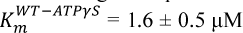 for TF_1_(WT), and 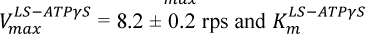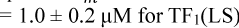. (C) Structural comparison of *b*MF_1_ (PDB: 2JDI) and TF_1_ (PDB: 8HH1), highlighting the bound nucleotide and the residue pairs L317/S336 in *b*MF_1_ and Y313/E332 in TF_1_. The β and γ subunits are shown in green and salmon pink, respectively. Dashed lines indicate the distances between the indicated residue pairs, which are 7.1 Å in *b*MF_1_ and 2.7 Å in TF_1_.

Previous kinetic analyses of TF_1_ rotation have established that the maximum rotation rate is primarily limited by ATP cleavage and phosphate release (30, 31). To determine whether the enhanced turnover resulted from faster ATP hydrolysis, we next analyzed rotation driven by ATPγS, a slowly hydrolyzable ATP analogue that specifically prolongs the ATP hydrolysis dwell in the rotation assay (21, 31) (Fig. 6B). If ATP hydrolysis were selectively accelerated in TF_1_(LS), ATPγS-driven rotation would also be expected to increase relative to TF_1_(WT). However, TF_1_(LS) instead showed a slightly lower maximum rotation rate than TF_1_(WT) under ATPγS-driven conditions. These results therefore suggest that the enhanced rotation of TF_1_(LS) results from acceleration of another rate-limiting step, such as phosphate release or a conformational transition associated with product release.

## Discussion

### Requirement for enzyme engineering

In this study, we increased the maximum rotation rate of TF_1_ by 1.8-fold while preserving its functional thermostability. Our engineering strategy integrated homolog-guided sequence comparison, targeted exploration of activity-enhancing hotspots, and machine-learning-assisted prioritization. This workflow efficiently navigated the sequence space of this complex molecular motor and identified mutations that would have been difficult to identify from current mechanistic understanding alone.

Homolog-guided protein engineering (32, 33) has been widely employed to improve protein thermostability and expression levels (34–38), whereas its application to activity enhancement has been relatively limited (39–41). One representative example is the engineering of thermophilic 3-isopropylmalate dehydrogenase (3-IPMDH), in which several substitutions derived from a mesophilic homolog markedly enhanced catalytic activity at 25°C without compromising thermostability (42). Compared with such soluble enzymes, F_1_-ATPase is a much larger multi-subunit molecular motor, in which ATP hydrolysis is tightly coupled to long-range conformational transitions. Such molecular complexity makes rational activity engineering considerably more challenging. Nevertheless, by introducing the *b*MF_1_–PdF_1_ consensus residues determined by comparative sequence analysis (Fig. 2A), we identified several activity-enhancing substitutions, particularly at residues distal to the catalytic site, demonstrating that homolog-guided sequence comparison provides an effective strategy for identifying functionally important hotspots even in a highly complex molecular motor.

Importantly, however, the optimal substitutions were not necessarily the *b*MF_1_–PdF_1_ consensus residues. For example, at βY313, several non-consensus substitutions increased the rotation rate to a similar extent as the *b*MF_1_–PdF_1_ consensus substitution. This behavior contrasts markedly with our previous saturation mutagenesis of first-shell catalytic residues, in which the native amino acids consistently exhibited the highest activities and most substitutions caused severe loss of function (24). Together, these observations suggest that first-shell catalytic residues are already highly optimized through evolution, whereas second-shell residues retain substantial evolutionary and engineering flexibility (Fig. S7). Homolog-guided substitutions therefore provide an effective strategy for identifying activity-enhancing hotspots within this designable sequence space, followed by focused exploration to identify the optimal amino acid at each hotspot.

Machine-learning-assisted prioritization further enriched highly active mutants within the experimentally accessible sequence space. Although the present model did not identify mutants superior to TF_1_(LS), most top-ranked candidates exhibited higher activities than TF_1_(WT), demonstrating that machine learning effectively reduced the experimental search space. The limited predictive performance of the model likely reflects the relatively small training dataset, as well as the strong epistatic interactions among mutations, as illustrated by the reduced activity of the quintuple mutant containing αR132K, αR383K, βY313L, βE332S, and the *b*-loop insertion (Fig. 4D). Future integration of larger experimental datasets with iterative model retraining may further improve prediction accuracy and facilitate exploration of broader combinatorial sequence spaces (27, 28, 43).

Our results also provide an example in which catalytic activity was enhanced without compromising functional thermostability. Enzymes generally require a balance between structural stability and conformational flexibility, and increased catalytic activity is often accompanied by reduced stability (44–54) . In contrast, TF_1_(LS) retained the functional thermostability of TF_1_(WT) while exhibiting higher catalytic activity. These findings support the idea that thermophilic proteins provide a stability margin that can accommodate activity-enhancing mutations without substantial functional destabilization (42, 55, 56). Such highly stable protein scaffolds, including proteins redesigned using computational protein design tools (57, 58), may therefore represent attractive starting points for future enzyme engineering.

### Possible molecular interpretation of the best mutant

The kinetic analysis (Fig. 6A and 6B) suggests that the enhanced rotation rate of TF_1_(LS) does not arise from accelerated ATP binding. Furthermore, the lower ATPγS-driven rotation rate of TF_1_(LS) argues against ATP cleavage being the primary mechanism responsible for the enhanced rotation. Instead, the major contribution is more likely to arise from accelerated phosphate release or a conformational transition associated with product release. This interpretation is consistent with the structural insights of the two substituted residues. βY313 and βE332 are located on the inward-facing side of the β subunit, approximately 10 Å from the bound nucleotide, and therefore represent second-shell rather than catalytic residues. Neither residue directly participates in ATP hydrolysis, suggesting that substantial enhancement of catalytic activity can be achieved without modifying the catalytic residues themselves. Instead, these substitutions are likely to alter local conformational dynamics surrounding the active site.

A structural comparison provides a possible mechanistic explanation for the enhanced activity of TF_1_(βY313L/βE332S) (Fig. 6C). In the available structures of TF_1_ (PDB: 8HH1) and *b*MF_1_ (PDB: 2JDI), the minimum side-chain heavy-atom distance is approximately 2.7 Å between βY313 and βE332 in TF_1_, but 7.1 Å between the corresponding βL317 and βS336 residues in *b*MF_1_. The shorter distance in TF_1_ may allow hydrogen-bond formation between βY313 and βE332, thereby increasing local structural rigidity. Replacement with βY313L and βE332S may weaken this interaction and facilitate conformational changes required for later catalytic events. Such subtle modulation of local flexibility may preferentially accelerate phosphate release or an associated conformational transition, thereby enhancing turnover rate. The contrasting effects of the LS substitutions on ATP- and ATPγS-driven rotation indicate that they do not enhance activity toward both substrates equally. This substrate-dependent behavior may reflect a possible constraint in engineering promiscuous enzymes (59, 60). Further structural analyses by cryo-EM, together with molecular dynamics simulations, will be valuable for testing the proposed mechanisms.

## Methods

### Determination of possible mutation sites from sequence alignments of three genuine F_1_

All possible mutation sites were identified through sequence alignment of the α- or β-subunit sequence of three genuine F_1_s: TF_1_, PdF_1_ and *b*MF_1_. The alignments were performed using the MEGA11 (Molecular Evolutionary Genetics Analysis version 12) (61) with the MUSCLE algorithm (62) under default settings. From the alignment, residues in which a TF_1_ residue differed from the PdF_1_-*b*MF_1_ consensus residues was defined as possible mutation sites. For the TF_1_(β^30^) mutant, all such residues located within the central part of the β-subunit (positions 150–413 in the TF_1_ β-subunit) were selected. For the TF_1_(*b*-loop) mutant, the mutation site corresponded to a gap observed in the alignment, where 6 residues present in *b*MF_1_ were inserted between positions 208 and 209 of the TF_1_ β-subunit (Fig. S2). Possible mutation sites within the α-subunit were determined in a similar way as for β-subunit, targeting on the α_DP_β_DP_ interface of TF_1_.

### Construction of mutant genes

For the plasmids encoding the α- and β-subunits used in the cell-free expression system, synthetic oligonucleotide fragments containing the mutation were purchased from Fasmac. The remaining plasmid regions, excluding the mutation-containing oligonucleotide fragment, were amplified by PCR using primers synthesized by Fasmac. The resulting DNA fragments were assembled using NEBuilder HiFi DNA Assembly to generate the mutant plasmids. The same procedure was also applied to generate TF_1_(*b*-loop) and TF_1_(βY313L/βE332S) plasmids for recombinant protein expression and purification in *E. coli*. For the TF_1_(β^30^) plasmid, a gene fragment encoding residues 150–413 with 30 mutations was codon-optimized for expression in *E. coli* and synthesized by Eurofins. The synthesized insert and the corresponding vector were digested with AgeI and PmlI, followed by ligation to generate the mutant plasmid. For point and combination mutants, experimental feasibility of oligonucleotide design, PCR amplification, and fragment assembly was considered when prioritizing constructs for screening.

### Single-molecule rotation assay of F_1_ with purified or cell-free expressed proteins

The purified proteins of TF_1_(WT), TF_1_(β^30^), TF_1_(*b*-loop), and TF_1_(βY313L/βE332S) were prepared as described in the previous paper (23). Single-molecule rotation assay of these purified proteins was performed as follows. The flow cell was constructed from two cover glasses (18 × 18 mm^2^ and 24 × 32 mm^2^; Matsunami Glass) using double-sided tape as a spacer. The surface of the bottom glass was coated with Ni-NTA. First, the flow cell was incubated with BSA buffer (the buffer containing 5 mg/mL BSA, 50 mM MOPS-KOH (pH 7.0), and 50 mM KCl, 2 mM MgCl_2_) for 5 min. Next, F_1_ molecules of 200 pM in the BSA buffer were infused and incubated for 5 min. Then, unbound F_1_ molecules were washed out by the BSA buffer and 40 nm gold nanoparticles were infused and incubated for 10 min. Unbound particles were washed out with the buffer for the assay containing 50 mM MOPS-KOH (pH 7.0), 50 mM KCl, 2 mM MgCl_2_, 100 μg/mL pyruvate kinase, 2.5 mM phosphoenolpyruvate, with the indicated ATP concentration. ATP-regenerating system (pyruvate kinase and phosphoenolpyruvate) was omitted in the assays using ATPγS. The assay was conducted with a dark-field microscope (OLYMPUS IX-71) at the recording rate of 1-10k fps (FASTCAM-1024PCI or FASTCAM NOVA s-16, Photron, Japan) (61). Movies were analyzed by a custom software (62). The localization precision and signal-to-noise ratio ranges were the same as the previous studies (21). Rotation rates were calculated from the elapsed time required for the fastest 50 consecutive revolutions. The rotation rates of all mutants examined in this study are summarized in Fig. S8.

TF_1_ mutants for high-throughput screening were expressed using a cell-free protein synthesis system as described previously (22) with some modifications. To avoid unexpected allosteric effect, the βT165S/G181A mutations were removed and the corresponding WT plasmid was used instead. The expression mixtures were diluted 50-fold with buffer containing 100 mM potassium phosphate (pH 7.0) and 2 mM EDTA, and then partially purified by centrifugation at 14,000 × g for 15 min using Amicon Ultra 100K centrifugal filter units (0.5 mL). The rotation assay for these mutants was conducted in a similar way to that for purified proteins, except for several modifications. The initial incubation with BSA buffer was omitted. Before sealing the flow cell with the top cover glass, 0.5–1 µL of each protein solution was directly applied onto the flow lane. After washing by BSA buffer, the lane was washed further with BSA buffer containing 50 mM imidazole–HCl (pH 7.2) to remove incomplete F_1_ complex and solution contaminants from cell-free expression. Subsequently, 40 nm gold nanoparticles were loaded, and the assay buffer was introduced to wash unbound gold nanoparticles. Observation was performed in the same manner as for the purified proteins.

### Machine learning

Measured rotation rates of 30 TF_1_ mutants containing mutations at one or more of the following sites were used as training data (Table. S1): βY313, βE332, αR132, αR383, and/or the *b*-loop insertion (Fig. 4). Mutants carrying any additional mutations outside these sites were excluded. The response variable was the experimentally measured rotation rate without transformation. Mutation information was encoded using amino-acid descriptor vectors, and the *b*-loop insertion was represented as a binary indicator. Eight descriptor families were considered (BLOSUM (63), FASGAI (64), MS-WHIM (65), ProtFP (66), ST-scale (67), T-scale (68), VHSE (69), and Z-scale (70)). Random Forest regressors were trained using scikit-learn (71), and the combination of T-scale, VHSE, and Z-scale, together with the model hyperparameters, was selected by five-fold cross-validation using Spearman’s rank correlation as the scoring metric (Fig. S4). Out-of-fold predictions from the five-fold cross-validation were used to assess rank-ordering performance. The final model was trained using this descriptor combination and the selected hyperparameters, and was then used to predict the rotation speeds of 320,000 mutants representing all possible amino-acid combinations at the four positions, in combination with the presence or absence of the *b*-loop insertion. Predicted values were ranked in descending order to identify candidate highly active mutants.

### Thermostability analysis of F_1_s

TF_1_(WT) and TF_1_(βY313L/βE332S) were purified as described above, and *b*MF_1_ was prepared as described in the reference article (72). All purified proteins were diluted to a final concentration of 3 µM in 100 mM potassium phosphate (pH 7.0) and 2 mM EDTA. Samples were preincubated at room temperature for over 10 minutes, then incubated in a thermal cycler either at 25 °C for 15 min or at 65 °C for 10 min followed by 25 °C for 5 min. Thermostability was evaluated as the ratio of residual ATPase activity after heat treatment (65 °C for 10 min followed by 25 °C for 5 min) to that after incubation at 25 °C for 15 min. ATPase activity was quantified from the NADH oxidation rate monitored at 340 nm (73). Measurements were performed at 25 °C on a UV/Vis spectrophotometer equipped with a temperature controller. The assay buffer contained 50 mM MOPS–KOH (pH 7.0), 50 mM KCl, 3 mM MgCl_2_, 2.5 mM phosphoenolpyruvate, 200 µg/mL pyruvate kinase, 50 µg/mL lactate dehydrogenase, 0.2 mM NADH, and 3 mM ATP. NADH was added during measurements when necessary. Reactions were initiated by adding the pre-equilibrated F_1_ to the assay mixture containing ATP.

### Structure prediction by AlphaFold3

The structure of the αβ complex was predicted by AlphaFold Server (74) with default settings. The β-subunit sequence in TF_1_ containing the *b*-loop insertion was used for the prediction. To obtain an ATP-bound model at the catalytic αβ interfaces, one α subunit and two β subunits were predicted with two ATP molecules. The resulting predicted structure was used to visualize the positions of the mutation hotspots in Fig. 3A.

## Supporting information

Supplemental infomation

## Acknowledgments

We thank all members of the Noji laboratory for their valuable comments. This work was supported in part by JSPS KAKENHI grants for Early-Career Scientists (Grant Number 25K18428 to R.K.) and Scientific Research on Innovative Areas (JP21H00388 to H.U.), Challenging Research (Exploratory) (JP23K18092 to H.U.), Scientific Research (B) (JP24K01987 to H.U.), and Scientific Research (S) (JP19H05624 to H.N.); a Research Grant from the Human Frontier Science Program (RGP0054/2020 to H.N.); and the JST ASPIRE Program (JPMJAP24B5 to H.N.).

## Author Contributions

R.K., K.M., H.U., and H.N. designed the research; R.K., K.M., T.O conducted experiments and analyses. R.K. and K.M constructed machine-learning with support from Y.S.. R.K., M.K. and H.N. wrote the paper with support from H.U. and Y.S.

## Data availability

The data that support the findings of this study are available from the corresponding author upon reasonable request.

## AI Disclosure Statement

During the preparation of this manuscript, the authors used ChatGPT (OpenAI) for drafting, editing, and rewriting manuscript sections; translation of content; summarization of manuscript sections and related text; synthesis and analysis of information from the literature; and generation or modification of code. AI-assisted output was used only as a supportive tool. The authors critically reviewed, edited, and verified all AI-assisted text, translations, summaries, literature-based analyses, and code, and take full responsibility for the accuracy, originality, and integrity of the final manuscript. AI technology was not used to create, alter, or manipulate original research data, research results, figures, images, or data visualizations.

