## Supplemental infomation for "Engineering a highly active thermophilic F_1_-ATPase by homolog-guided exploration and machine-learning-assisted prioritization"

This PDF file includes:

- Figs. S1 to S8
- Table. S1

[1] *Department of Applied Chemistry, Graduate School of Engineering, The University of Tokyo, 7-3-1 Hongo, Bunkyo-ku, Tokyo 113-8656, Japan*
[2] *Artificial Intelligence Research Center, National Institute of Advanced Industrial Science and Technology (AIST), 2-4-7 Aomi, Koto-ku, Tokyo 135-0064, Japan*
[3] *Department of Computational Biology and Medical Sciences, Graduate School of Frontier Sciences, The University of Tokyo, 5-1-5 Kashiwanoha, Kashiwa, Chiba 277-8563, Japan*
[4] *Department of Data Science, School of Frontier Engineering, Kitasato University, 1-15-1 Kitasato, Minami-ku, Sagamihara, Kanagawa 252-0373, Japan*
[5] *Research Institute of Planetary Health (RIPH), The University of Tokyo, 2-21-2 Takanawa, Minato-ku, Tokyo 108-0074, Japan*

[†] *These authors contributed equally*

* Corresponding author: Hiroyuki Noji

```
TF1  MTR-----------GRVIQVMGPVVDVKFENGHLPAIYNALKIQHKARNENEVDIDLTLEVALHLGDDTVRTIAMASTDGLIR
bMF1 AAQASPSPKAGATTGRIVAVIGAVVDVQFDEG-LPPILNALEVQGRETR-------LVLEVAQHLGESTVRTIAMDGTEGLVR
PdF1 MAEAN---------GKITQVIGAVVDVQFD-GQLPAILNALETENNGKR-------LVLEVAQHLGENTVRTIAMDATEGLVR

                                                                N-terminus was removed
     GMEVIDTGAPISVPVGEVTLGRVFNVLGEPIDLEGDIPADARRDPIHRPAPKFEELATEVEILETGIKVVDLLAPYIKGGKIG
     GQKVLDSGAPIRIPVGPETLGRIMNVIGEPIDERGPIKTKQFA-AIHAEAPEFVEMSVEQEILVTGIKVVDLLAPYAKGGKIG
     GLPVKDTGGPIMVPVGDATLGRILNVVGEPVDEGGPVEATQTR-AIHQQAPDFAAQATASEILVTGIKVIDLLAPYSKGGKIG

       P-loop           Hinge   General base           Insertion (b-loop)
     LFGGAGVGKTVLIQELIHNIAQEHGGISVFAGVGERTREGNDLYHEMKDSGVI-------SKTAMVFGQMNEPPGARMRVALT
     LFGGAGVGKTVLIMELINNVAKAHGGYSVFAGVGERTREGNDLYHEMIESGVINLKD-ATSKVALVYGQMNEPPGARARVALT
     LFGGAGVGKTVLIMELINNIAKVHSGYSVFAGVGERTREGNDLYHEMVESGVIKPDDLSKSQVALVYGQMNEPPGARMRVALT

                                                                                    Y313
     GLTMAEYFRDEQGQDVLLFIDNIFRFTQAGSEVSALLGRMPSAVGYQPTLATEMGQLQERITSTAKGSITSIQAIYVPADDYT
     GLTVAEYFRDQEGQDVLLFIDNIFRFTQAGSEVSALLGRIPSAVGYQPTLATDMGTMQERITTTKKGSITSVQAIYVPADDLT
     GLTVAEQFRDATGTDVLFFVDNIFRFTQAGSEVSALLGRIPSAVGYQPTLATDMGAMQERITSTKNGSITSIQAVYVPADDLT

                  E332                                                       DELSEED loop
     DPAPATTFSHLDATTNLERKLAEMGIYPAVDPLASTSRALAPEIVGEEHYQVARKVQQTLQRYKELQDIIAILGMDELSDEDK
     DPAPATTFAHLDATTVLSRAIAELGIYPAVDPLDSTSRIMDPNIVGSEHYDVARGVQKILQDYKSLQDIIAILGMDELSEEDK
     DPAPATTFAHLDATTVLSRAISELGIYPAVDPLDSNSRILDPAVVGEEHYQVARDVQGILQKYKSLQDIIAILGMDELSEEDK

                      C-terminus was removed
     LVVHRARRIQFFLSQNFHVAEQFTGQPGSYVPVKETVRGFKEILEGKYDHLPEDAFRLVGRIEEVVEKAKAMGVEV-
     LTVSRARKIQRFLSQPFQVAEVFTGHLGKLVPLKETIKGFQQILAGEYDHLPEQAFYMVGPIEEAVAKADKLAEEHS
     LTVARARKIQRFLSQPFDVAKVFTGSDGVQVPLEDTIKSFKAVVAGEYDHLPEAAFYMVGGIEDVKAKAQRLAADAA
```

**Fig. S1. Sequence alignment of the β subunit**

The sequences of the β subunit from $TF_1$ (accession number: P07677), *b*$MF_1$ (accession number: P00829), and $PdF_1$ (accession number: A1B8P0) were analyzed by MEGA11. The amino acid residues written in bold blue and red represent the candidate mutation sites and substitutions employed in $TF_1(\beta^{30})$.

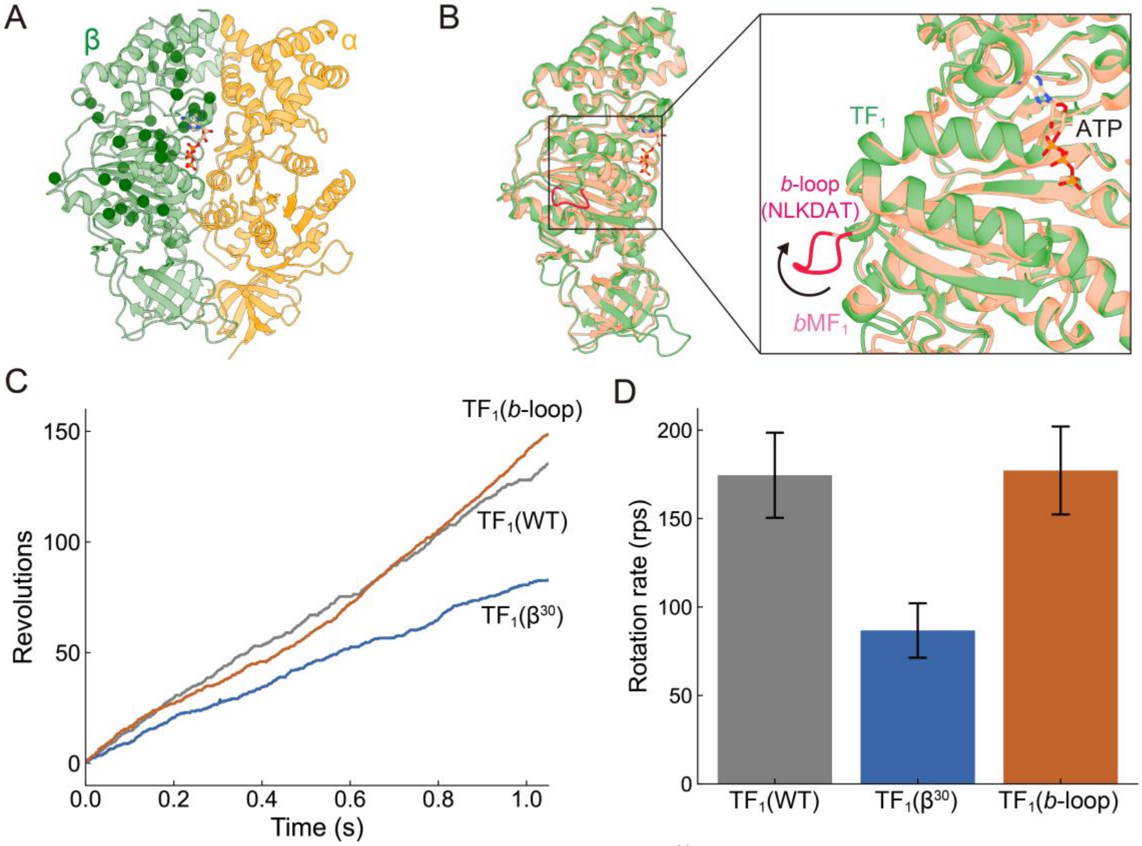


**Fig. S2. Single-molecule rotation assay of purified $TF_1(\beta^{30})$ and $TF_1$(*b*-loop).**
(A) Mutation sites of $TF_1(\beta^{30})$ plotted onto the $\alpha\beta_{DP}$ state in the $TF_1$ structure (PDB: 7l1r), showing mutation sites as a sphere shape. $TF_1(\beta^{30})$ includes the following 30 mutations: Q169M, H173N, Q177K, I182Y, D204E, T211V, M213L, F215Y, M235V, M271I, E284D, L288M, A296K, Y313L, S323A, N330V, E332S, K334A, L335I, M338L, A348D, A353I, A355D, T373I, E379S, D394E, V399T, R405K, F408R, N413P. (B) Structural comparison of the $\beta_{DP}$ subunits in $TF_1$ structure (PDB: 7l1r, green) and *b*$MF_1$ (PDB: 2jdi, light pink), emphasizing the additional loop region in *b*$MF_1$, referred to as $TF_1$(*b*-loop) (deep pink). The arrow indicates the N-to-C direction. The ATP molecule shown here is located at the catalytic site in the $TF_1$ structure. Sequence alignment of the β subunit is shown in Fig. S1. (C) Representative rotary trajectories and x-y plots of $TF_1$(WT), $TF_1(\beta^{30})$, and $TF_1$(*b*-loop) at 3 mM ATP. (D) Average rotation rates of $TF_1$(WT), $TF_1(\beta^{30})$, and $TF_1$(*b*-loop) at 3 mM ATP. Bars and error bars represent the mean and SD from more than three independent molecules.

```
TF1   MSIRAEEISALIKQQIENYESQIQVSDVGTVIQVGDGIARAHGLDNVMSGEAVEFANAVMGMALNLEENNVGIVILGPYTGIKEGDEV
bMF1  QKTGTAEVSSILEERILGADTSVDLEETGRVLSIGDGIARVHGLRNVQAEEMVEFSSGLKGMSLNLEPDNVGVVVFGNDKLIKEGDIV
PdF1  MGIQAAEISAILKDQIKNFGQDAEVAEVGQVLSVGDGIARVYGLDKVQAGEMVEFPGGIRGMVLNLETDNVGVVIFGDDRDIKEGDTV

                                                   R132
      RRTGRIMEVPVGETLIGRVVNPLGQPVDGLGPVETTETRPIESRAPGVMDRRSVHEPLQTGIKAIDALVPIGRGQRELIIGDRQTGKT
      KRTGAIVDVPVGEELLGRVVDALGNAIDGKGPIGSKARRRVGLKAPGIIPRISVREPMQTGIKAVDSLVPIGRGQRELIIGDRQTGKT
      KRTGAIVEVPAGKELLGRVVDALGNPIDGKGPLNASERRIADVKAPGIMPRKSVHEPMATGLKSVDAMIPVGRGQRELIIGDRQTGKT

      SVAIDTIINQKD---------QNMICIYVAIGQKESTVATVVETLAKHGAPDYTIVVTASASQPAPLLFLAPYAGVAMGEYFMIMGKH
      SIAIDTIINQKRFND-GTDEKKKLYCIYVAIGQKRSTVAQLVKRLTDADAMKYTIVVSATASDAAPLQYLAPYSGCSMGEYFRDNGKH
      AIALDTILNQANYNGREADGMKTLHCIYVAVGQKRSTVAQLVKKLEETGAMAYTTVVAATASDPAPMQYLAPYSATAMGEYFRDNGMD

      VLVVIDDLSKQAAAYRQLSLLLRRPPGREAYPGDIFYLHSRLLERAAKLSDAKGGGSLTALPFVETQAGDISAYIPTNVISITDGQIF
      ALIIYDDLSKQAVAYRQMSLLLRRPPGREAYPGDVFYLHSRLLERAAKMNDAFGGGSLTALPVIETQAGDVSAYIPTNVISITDGQIF
      ALIIYDDLSKQAVAYRQMSLLLRRPPGREAYPGDVFYLHSRLLERSAKLNEANGAGSLTALPIIETQAGDVSAYIPTNVISITDGQIF

                   Arginine Finger          R383
      LQSDLFFSGVRPAINAGLSVSRVGGAAQIKAMKKVAGTLRLDLAAYRELEAFAQFGSDLDKATQANVARGARTVEVLKQDLHQPIPVE
      LETELFYKGIRPAINVGLSVSRVGSAAQTRAMKQVAGTMKLELAQYREVAAFAQFGSDLDAATQQLLSRGVRLTELLKQGQYSPMAIE
      LETELFFQGIRPAVNTGLSVSRVGSAAQTKAMKSVAGPVKLELAQYREMAAFAQFGSDLDAATQKLLNRGARLTELMKQPQYSPLTNA

      KQVLIIYALTRGFLDDIPVEDVRRFEKEFYLWLDQNGQHLLEHI-RTTKDLPNE--DDLNQAIEAFKKTFVVSQ
      EQVAVIYAGVRGYLDKLEPSKITKFENAFLSHVISQHQALLSKI-RTDGKISEESDAKLKEIVTNFLAGFEA--
      EIVIVIYAGTKGYLDGIPVRDVTKWEHGLLQYLRNQKADLLEDMTKNDRKVAGELEDAIKAALDGYAKTYA---
```

**Fig. S3. Sequence alignment of the α subunit**

The sequences of the α subunit from $TF_1$ (accession number: P09219), *b*$MF_1$ (accession number: P19483), and $PdF_1$ (accession number: A1B8N8) were analyzed by MEGA11.

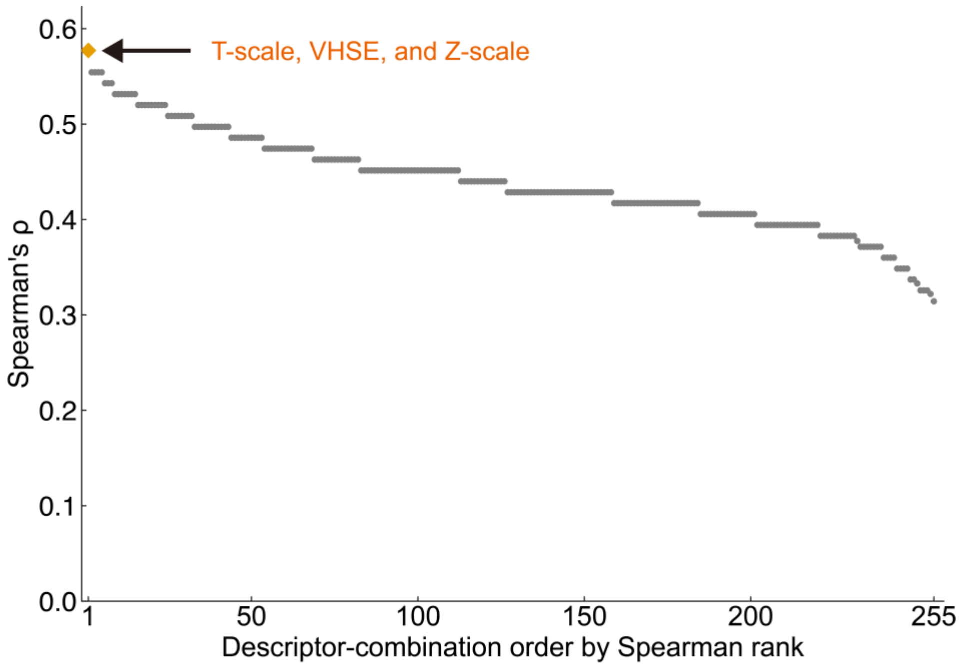


**Fig. S4. Comparison of amino-acid descriptor combinations based on Spearman's rank correlation coefficient (ρ).**

The 255 combinations are ordered from highest to lowest ρ. The orange diamond indicates the best-performing combination, comprising T-scale, VHSE, and Z-scale, and gray circles indicate all other combinations.

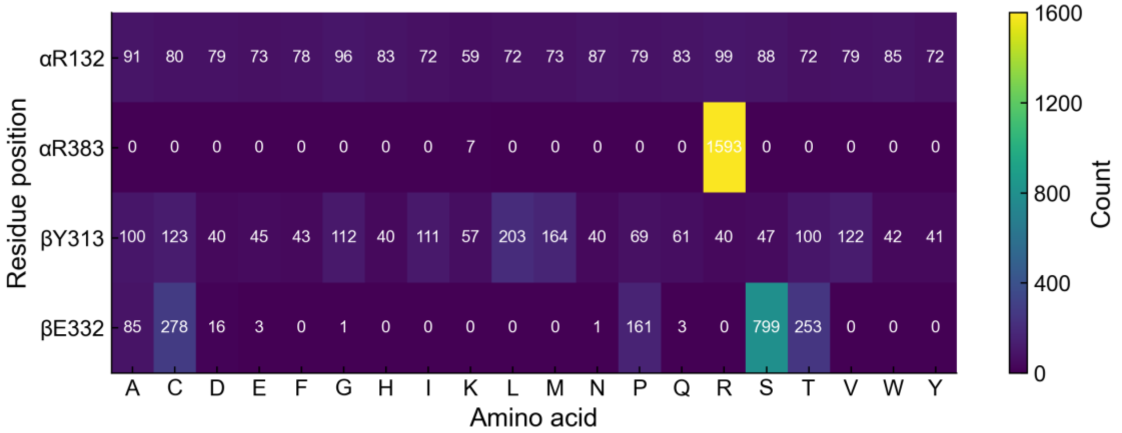


**Fig. S5. Amino-acid distribution among the top-ranked predicted mutants.**

The heatmap shows the amino acid composition at four substituted residue positions (αR132, αR383, βY313, βE332) among the top 0.5% (1600) ranked mutants. Values in each cell represent the number of mutants containing the corresponding amino acid at that position. The color scale indicates the occurrence count.

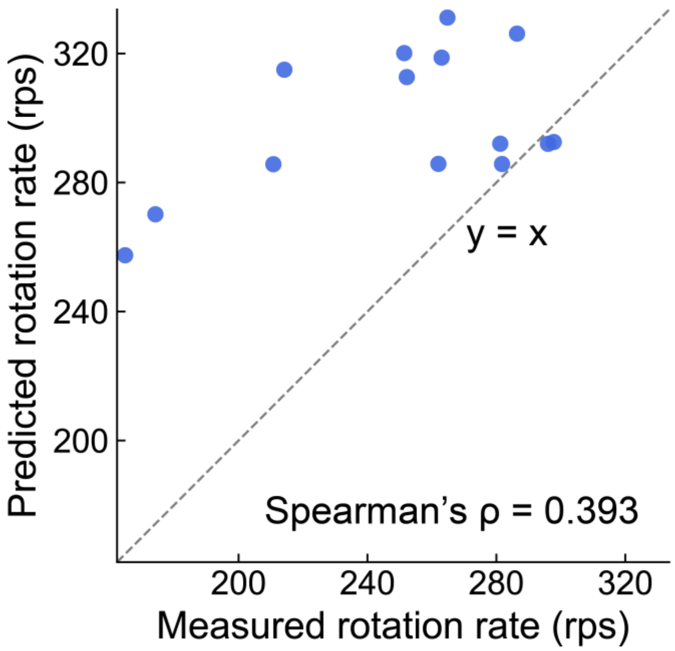


**Fig. S6. Relationship between measured and predicted rotation rates of mutants.**

Each point represents an individual mutant. The gray dashed line represents the 1:1 line, where predicted values equal measured values. Most points were located above the 1:1 line, indicating that the model tended to overestimate the rotation rate.

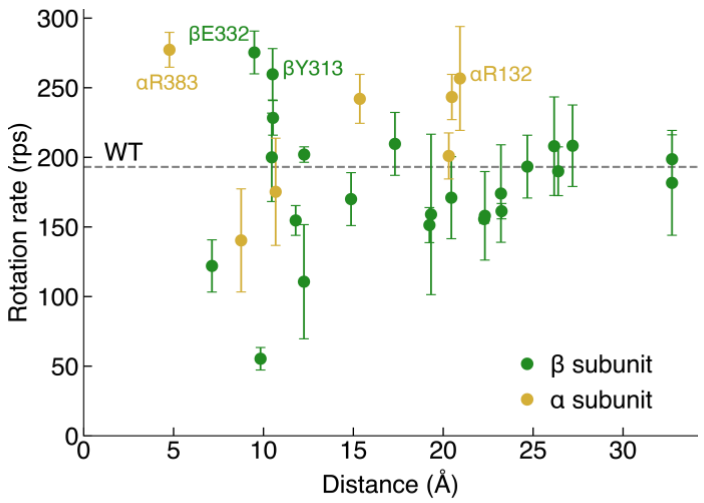


**Fig. S7. Relationship between the minimum ATP–residue distance and rotation rate.**
The rotation rate is plotted against the shortest atom-to-atom distance between each residue and the ATP molecule at the $\alpha\beta_{DP}$ site in the 7L1R structure. The residues in the α and β subunit are shown in yellow and green, respectively. Mutants containing multiple substitutions were excluded, and only point mutations are shown. Error bars represent the SD from at least three independent molecules. The gray dashed line indicates the rotation rate of WT, 193 rps.

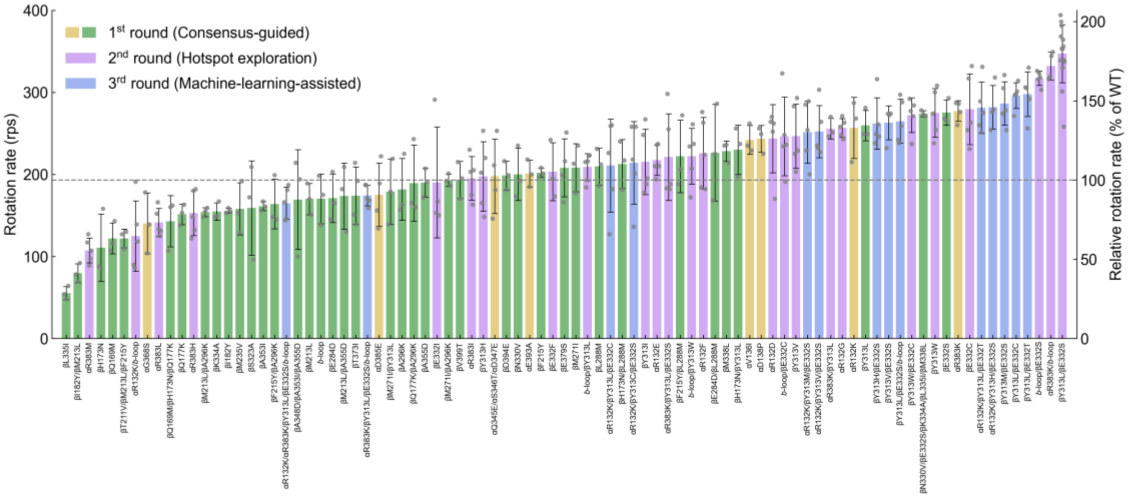


**Fig. S8. Rotation rates of $TF_1$ mutants tested in this study.**

Bar colors indicate each screening round. Bars and error bars represent the mean ± SD for at least three independent molecules, and gray dots indicate rotation rates from individual molecules. The dashed horizontal line indicates the mean rotation rate of $TF_1$(WT) (193 rps). The right y-axis shows rotation rates relative to that of $TF_1$(WT).

**Table. S1. Experimental dataset used for machine-learning-assisted exploration.**
The dataset contains the experimentally measured rotation rates of 30 $TF_1$ mutants carrying substitutions at βY313, βE332, αR132, αR383, and with or without the *b*-loop insertion.

| Mutations | Average rotation rates (rps) |
|---|---|
| WT | 193.1 |
| αR132D | 243.6 |
| αR132E | 217.4 |
| αR132F | 225.8 |
| αR132G | 256.6 |
| αR132K | 256.7 |
| αR383I | 195.2 |
| αR383L | 141.4 |
| αR383M | 107.4 |
| αR383H | 152.8 |
| αR383K | 277.3 |
| βY313H | 197.6 |
| βY313I | 215.2 |
| βY313V | 246.6 |
| βY313W | 274.8 |
| βY313L | 259.7 |
| βE332C | 279.4 |
| βE332F | 203.3 |
| βE332I | 190.3 |
| βE332S | 275.3 |
| αR383K/βY313L | 255.7 |
| αR383K/*b*-loop | 332.0 |
| αR132K/*b*-loop | 124.8 |
| βY313W/*b*-loop | 222.2 |
| βY313L/*b*-loop | 209.0 |
| βE332C/*b*-loop | 246.4 |
| βE332S/*b*-loop | 317.2 |
| βY313W/βE332C | 272.2 |
| βY313L/βE332S | 347.1 |
| αR383K/βY313L/βE332S | 221.2 |
